# Phosphatidylinositol 3,5-Bisphosphate regulates Ca^2+^ Transport During Yeast Vacuolar Fusion through Activation of the Ca^2+^ ATPase Pmc1

**DOI:** 10.1101/533067

**Authors:** Gregory E. Miner, Katherine D. Sullivan, Annie Guo, Matthew L. Starr, EZ C. Ellis, Brandon C. Jones, Rutilio A. Fratti

## Abstract

The transport of Ca^2+^ across membranes precedes the fusion and fission of various lipid bilayers. Yeast vacuoles during hyperosmotic shock become fragmented through fission events that require Ca^2+^ efflux of their luminal stores through the TRP channel Yvc1. This requires the production of the lipid PI(3,5)P_2_ by Fab1. Ca^2+^ is also released during vacuole fusion upon *trans*-SNARE complex assembly, however, the role of PI(3,5)P_2_ remains unclear. Here we demonstrate that elevated PI(3,5)P_2_ levels abolish Ca^2+^ efflux during fusion, indicating that PI(3,5)P_2_ has opposing effects on Ca^2+^ transport in fission versus fusion. Notably, Ca^2+^ efflux was enhanced when PI(3,5)P_2_ levels were reduced. Importantly, the effect of PI(3,5)P_2_ on Ca^2+^ flux was independent of Yvc1. Rather, the effect was dependent on the Ca^2+^ pump Pmc1. Vacuoles lacking Pmc1 were resistant to the effects of PI(3,5)P_2_, while those lacking Yvc1 remained sensitive. Furthermore altering PI(3,5)P_2_ levels affects the interactions of Pmc1 with the V_o_ component Vph1 and the R-SNARE Nyv1. We now propose a model in which elevated PI(3,5)P_2_ activates continued Pmc1 function to prevent the accumulation of released extraluminal Ca^2+^.

**Summary:** During osmotic stress PI(3,5)P_2_ triggers Ca^2+^ release from vacuoles. Here we show PI(3,5)P_2_ stimulates Ca^2+^ uptake by vacuoles during fusion, illustrating that it has a dual role in Ca^2+^ transport.

## INTRODUCTION

The role of Ca^2+^ as a regulator of membrane fusion is best understood in the release of neurotransmitters through the fusion of synaptic vesicles with the presynaptic plasma membrane (Südhof, 2012). Upon plasma membrane depolarization an influx of Ca^2+^ into the cytoplasm occurs which subsequently binds to the C2 domains of synaptotagmin-1. Once bound to Ca^2+^, synaptotagmin-1 changes conformation and aids in triggering SNARE-mediated fusion (Chapman, 2008). Ca^2+^ transport across membranes is also a part of endolysosomal trafficking. In *Saccharomyces cerevisiae* vacuolar lysosomes serve as the major Ca^2+^ reservoir where it is complexed with inorganic polyphosphate (Dunn et al., 1994). The transport of Ca^2+^ into the vacuole lumen is mediated through the Ca^2+^/H^+^ exchanger Vcx1, and the ATP driven pump Pmc1, whereas the movement out of the lumen into the cytoplasm is driven by the mucolipin transient receptor potential cation channel (TRPML) family orthologue Ca^2+^ channel Yvc1 (Cunningham, 2011).

Although incomplete, we have a partial understanding of the mechanisms driving the release of Ca^2+^ during intracellular membrane trafficking. During osmotic shock or mechanical stress Yvc1 transports Ca^2+^ from the vacuole lumen to the cytosol that is controlled in part by the formation of PI(3,5)P_2_ by the PI3P 5-kinase Fab1/PIKfyve (Bonangelino et al., 2002; Denis and Cyert, 2002; Dong et al., 2010; Su et al., 2011). Through unresolved mechanisms this cascade leads to the fragmentation of the vacuole into small vesicles through fission events.

During membrane fusion the release of Ca^2+^ is triggered by *trans*-SNARE pairing (Merz and Wickner, 2004). However, a role for PI(3,5)P_2_ for fusion related Ca^2+^ efflux has not been tested. Separately, we showed that vacuole fusion was inhibited in the presence of elevated PI(3,5)P_2_ (Miner et al., 2019). Fusion was blocked by either the addition of an exogenous dioctanoyl form of the lipid (C8-PI(3,5)P_2_) or through endogenous overproduction by the hyperactive *fab1*^T2250A^ mutant. Fusion was arrested after the formation of *trans*-SNARE pairs but before mixing of the outer leaflets, i.e. hemifusion. The possible role for the TRP Ca^2+^ channel Yvc1 in the inhibition of fusion by PI(3,5)P_2_ was eliminated, as vacuoles lacking the transporter fused well and were equally sensitive to PI(3,5)P_2_ relative to wild type vacuoles. Although Yvc1 was not involved in the arrest of fusion, others have shown that Ca^2+^ efflux still occurs in the absence of the channel (Merz and Wickner, 2004). Thus, we postulated that PI(3,5)P_2_ could still affect Ca^2+^ transport through another mechanism.

Various mechanisms control the involvement of Ca^2+^ transport in membrane fusion including Ca^2+^-binding protein calmodulin and a specific ABC transporter. Calmodulin acts as a sensor and is participates in triggering late fusion events. Calmodulin tightly binding vacuoles upon Ca^2+^ release signals the completion of docking, which stimulates bilayer mixing (Ostrowicz et al., 2008; Peters and Mayer, 1998). Another mechanism that regulates fusion events through modulating Ca^2+^ transport involves the vacuolar Class-C ABC transporters Ybt1 (Sasser et al., 2012b; Sasser and Fratti, 2014).

Here we continued our study of PI(3,5)P_2_ during fusion and found that elevated concentrations of the lipid blocked the release of Ca^2+^. Both exogenous C8-PI(3,5)P_2_ and overproduction by *fab1*^T2250A^ inhibited Ca^2+^ efflux. In contrast, Ca^2+^ efflux was enhanced when Fab1 was inhibited by Apilimod, or when PI(3,5)P_2_ was sequestered by ML1-N. This effect was reproduced by the kinase dead *fab1^EEE^* mutation. Finally, we found that the effect of PI(3,5)P_2_ on Ca^2+^ transport occurs through its action on the Ca^2+^ uptake pump Pmc1 and not Yvc1. Thus, we conclude that the observed Ca^2+^ efflux during vacuole homotypic fusion is due in part to the regulation of Ca^2+^ influx by Pmc1.

## RESULTS

### Ca^2+^ Efflux is inhibited by PI(3,5)P_2_

The release of vacuole luminal Ca^2+^ stores occurs upon *trans*-SNARE complex formation in the path towards fusion (Merz and Wickner, 2004). The formation of vacuolar SNARE complexes also relies on the assembly of membrane microdomains enriched in regulatory lipids that include phosphoinositides such as the Fab1 substrate PI3P (Fratti et al., 2004). However, the link between phosphoinositides and Ca^2+^ transport during fusion has remained unclear. By contrast, the dependence of Ca^2+^ efflux on PI(3,5)P_2_ during vacuole fission under hyperosmotic conditions has been established (Dong et al., 2010). Together, this served as the impetus to explore the role of PI(3,5)P_2_ in fusion-linked Ca^2+^ transport.

To address this gap in knowledge, we used isolated vacuoles for *in vitro* assays of Ca^2+^ flux. Ca^2+^ transport was monitored through changes in Fluo-4 (or Cal-520) fluorescence as previously demonstrated (Miner et al., 2016; Miner et al., 2017; Miner and Fratti, 2019; Sasser et al., 2012b). Vacuoles were incubated under fusion conditions and reactions were started by the addition of ATP. As seen before, the addition of ATP triggers the rapid uptake of extraluminal Ca^2+^ from the buffer into the vacuole lumen as shown by the decrease in fluorescence (Fig. 1A, black line). Omission of ATP prevents Ca^2+^ uptake and fluorescence remained stable (not shown). Shortly after, and in a SNARE-dependent manner, Ca^2+^ is released from vacuoles as shown by an increase in fluorescence. As a negative control for SNARE-dependent efflux, select reactions were treated with antibody against Sec17 (*α*-SNAP), a co-chaperone of Sec18 (NSF) that aids in disrupting inactive *cis*-SNARE complexes to initiate the fusion pathway (Fig. 1A, dotted line) (Mayer et al., 1996).

**Figure 1.**
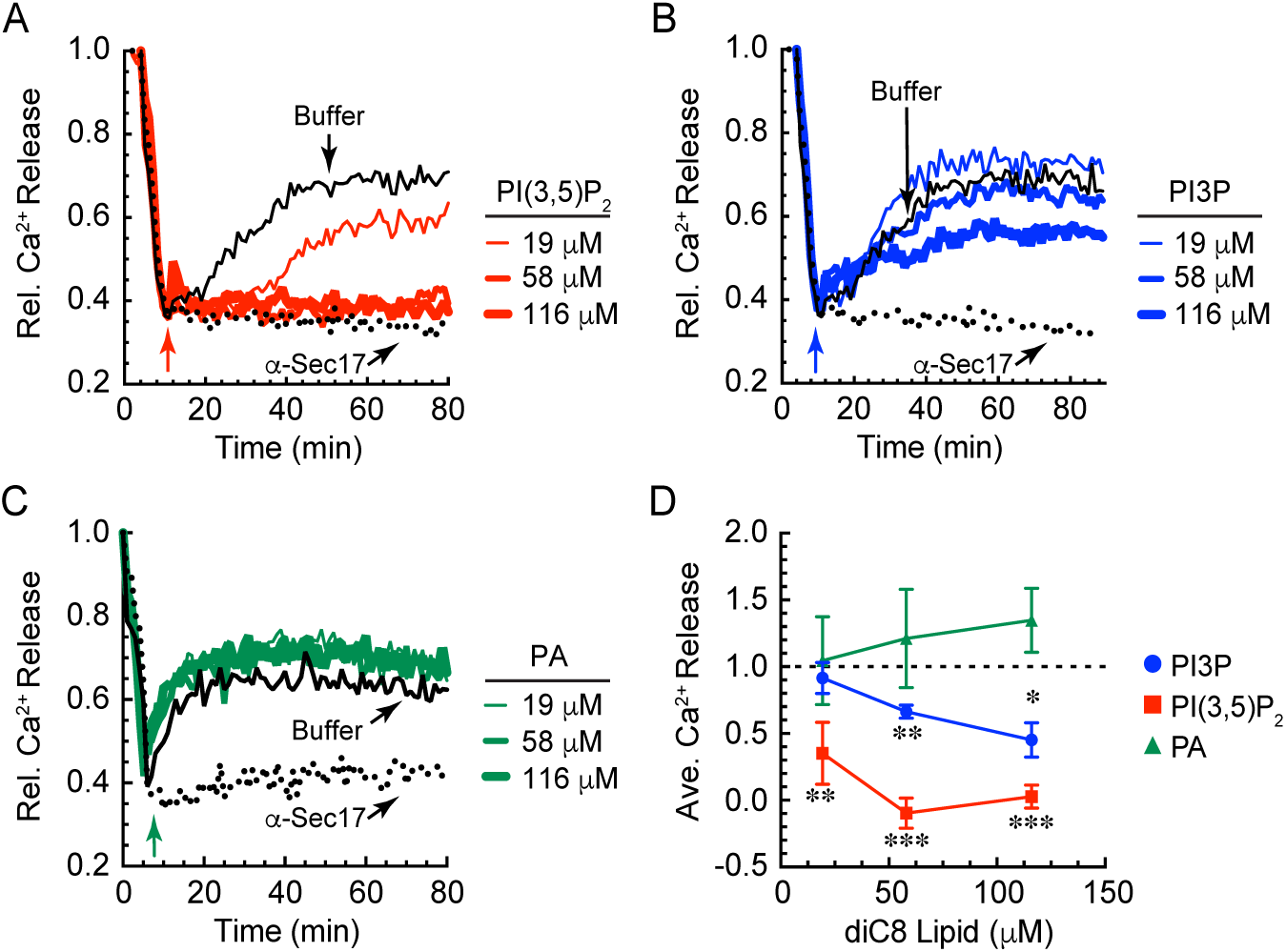
PI(3,5)P_2_ blocks Ca^2+^ efflux from vacuoles during fusion. Vacuoles were harvested from BJ3505 and 2X fusion reactions containing 150 µM Fluo-4 dextran. After 10 min of incubation with ATP (arrow) at 27°C to allow for the uptake of Ca^2+^, reactions were treated with C8-PI(3,5)P_2_ (A), C8-PI3P (B), or C8-PA (C) at the indicated concentrations. Reactions were further incubated at 27°C and fluorescence was measured every 30 sec for 80 min. Values were normalized to 1.0 representing the extraluminal Ca^2+^ at the beginning of the reaction and expressed as relative (Rel.) values compared to the untreated control. Separate reactions were incubated with 85 μg/ml anti-Sec17 IgG to block SNARE-dependent Ca^2+^ efflux. (D) Average Ca^2+^ efflux from A-C. Error bars are S.E.M. (n=3). \**p*<0.05, \*\**p*<0.01, \*\*\**p*<0.001.

Here we tested the effects of PI(3,5)P_2_ on Ca^2+^ transport. We treated reactions with dioctanoyl (C8) derivatives of PI(3,5)P_2_, as well as PI3P and phosphatidic acid (PA). PA was included due to its inhibitory effect on vacuole fusion (Starr et al., 2016), while PI3P was included as a precursor to PI(3,5)P_2_ and as a pro-fusion lipid (Boeddinghaus et al., 2002; Karunakaran et al., 2012; Karunakaran and Fratti, 2013; Lawrence et al., 2014; Miner et al., 2016). Short chain lipids were used to avoid solubility issues. As a result, seemingly high concentrations of C8 lipids were needed to see an effect. This, however, does not reflect the mol% of the lipids that partition into the membrane bilayer. Collins and Gordon showed that only a small fraction of C8-lipids incorporate into the membrane (Collins and Gordon, 2013). Based on their work we estimate that 100 µM C8-PI(3,5)P_2_ correlates to ∼1.2 mol% partitioning to the membrane fraction, which is in keeping with what reconstituted proteoliposome studies use in fusion models (Mima and Wickner, 2009). Lipids were added after 10 min of incubation to allow uniform uptake of Ca^2+^. We found that C8-PI(3,5)P_2_ blocked Ca^2+^ efflux in a dose dependent manner and as effectively as the *α*-Sec17 IgG negative control (Fig. 1A, D, red lines). In parallel we found that C8-PI3P only showed a modest inhibition at the highest concentration tested when compared to C8-PI(3,5)P_2_ (Fig. 1B, D). When C8-PA was tested we found that it did not significantly affect Ca^2+^ transport (Fig. 1C, D). These data indicate that the Ca^2+^ efflux associated with vacuole fusion was blocked by PI(3,5)P_2_. This is in contrast to the stimulatory effect of PI(3,5)P_2_ on Ca^2+^ efflux associated with vacuole fission. Importantly the concentrations used do not fully inhibit fusion, suggesting that Ca^2+^ transport can be uncoupled from fusion by altering the lipid composition of vacuole (Miner et al., 2019).

### Inhibition of the PI3P 5-Kinase Fab1/PIKfyve leads to enhanced Ca^2+^ Efflux

To verify the effect of PI(3,5)P_2_ on Ca^2+^ efflux we sought to counter its effects through inhibiting its production. The small molecule Apilimod has been shown to inhibit mammalian PIKfyve (Cai et al., 2013; Dayam et al., 2015). Thus, we first tested if Apilimod could inhibit yeast Fab1 activity. For this we measured the conversion of BODIPY-TMR C6-PI3P to BODIPY-TMR C6-PI(3,5)P_2_ by isolated vacuoles. Before we tested the effects of Apilimod, we verified our detection system by using *fig4Δ* and *fab1Δ* vacuoles that reduce or abolish PI(3,5)P2 production, respectively. In Figure 2A-B, we show that wild type vacuoles phosphorylated BODIPY-TMR C6-PI3P to make BODIPY-TMR C6-PI(3,5)P_2_. The half-life of the product was observed to be short as the levels of BODIPY-TMR C6-PI(3,5)P_2_ was significantly reduced after 5 min of incubation. This was in keeping reports by others showing that PI(3,5)P_2_ has a short half-life during vacuole fission (Duex et al., 2006a; Duex et al., 2006b; Jin et al., 2008). Next, vacuoles were incubated in the presence of BODIPY-TMR C6-PI3P and treated with either vehicle or Apilimod to inhibit Fab1 activity. Indeed, Apilimod treatment abolished BODIPY-TMR C6-PI(3,5)P_2_ generation (Fig. 2C-D).

**Figure 2.**
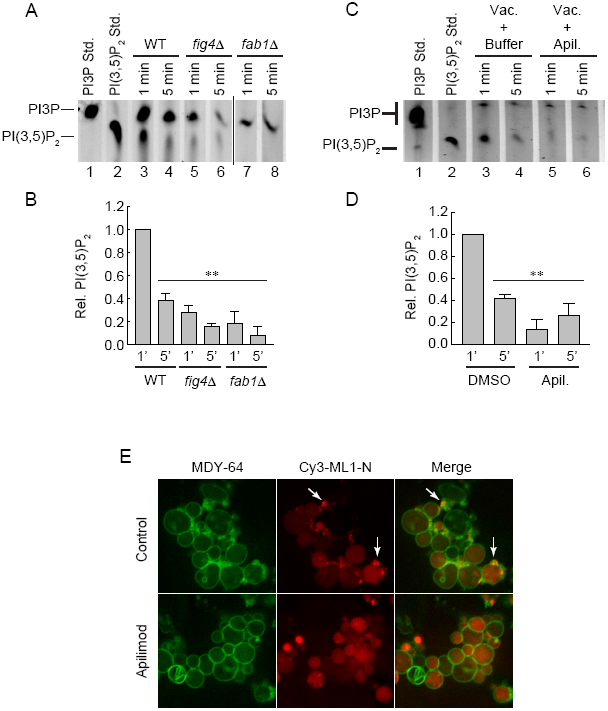
Apilimod inhibits Fab1 activity. (A) Vacuoles were incubated with BODIPY-TMR C6-PI3P at 27°C after which reactions were quenched with acetone and lipids were extracted and resolved by TLC to detect the production of BODIPY-TMR C6-PI(3,5)P_2_ by Fab1 activity. Wild type vacuoles were tested in parallel to those from *fig4Δ* and *fab1Δ* strains. Pure BODIPY-TMR C6-PI3P and C6-PI(3,5)P_2_ were used as standards in lanes 1 and 2. (B) Average of three experiments in panel A. (C) Vacuoles were treated with DMSO (carrier) or 500 µM Apilimod. (D) Average of three experiments in panel C. (E) Vacuoles were incubated under docking conditions for 20 min. Endogenous PI(3,5)P_2_ was labeled with 2.5 µM Cy3-ML1-N and the total membranes were labeled with 1 µM MDY-64. Vacuoles were treated with Apilimod or DMSO. Arrows point at examples of PI(3,5)P_2_ puncta. Error bars are S.E.M. (n=3).

To visualize the effect of Apilimod on intact vacuoles we performed docking assays where endogenously produced PI(3,5)P_2_ was labeled with Cy3-ML1-N as described previously (Miner et al., 2019). ML1-N is an N-terminal polypeptide from the endolysosomal mucolipin transient receptor potential (TRPML) Ca^2+^ channel (Dong et al., 2010). ML1-N preferentially binds to PI(3,5)P_2_ on isolated yeast vacuoles as no labeling is observed with vacuoles from the kinase inactive *fab1^EEE^* mutant (Miner et al., 2019). In Figure 2E we show that vacuoles treated with carrier (DMSO) showed distinct PI(3,5)P_2_ punctate staining as previously reported. When vacuoles were treated with Apilimod, we found that Cy3-ML1-N puncta were eliminated, further illustrating that Fab1 activity was inhibited during the experiment. Some free dye accumulated in the vacuole lumen to give the background fluorescence.

### Apilimod alters Ca^2+^ transport

The effects of Apilimod were next tested on Ca^2+^ efflux. We added a dose response curve of Apilimod after Ca^2+^ was taken up by vacuoles. The addition of Apilimod enhanced the release of Ca^2+^ in a dose dependent manner (Fig. 3A-B). This suggests that blocking Fab1 production of PI(3,5)P_2_ promotes Ca^2+^ efflux, which compliments the inhibition of Ca^2+^ release through the addition of C8-PI(3,5)P_2_ shown in Figure 1. The DMSO carrier control had no effect as shown by the overlapping curve with buffer control. It is important to note that pre-existing PI(3,5)P_2_ remained on the vacuole upon Apilimod treatment. To compare the effects of new lipid production with the starting pool of PI(3,5)P_2_ we tested whether physically blocking PI(3,5)P_2_ with ML1-N would reproduce the effects seen with Apilimod. When ML1-N was added we found that efflux was increased, which partially reproduced the Apilimod affect (Fig. 3C-D). However, higher concentrations (≥ 1 μM) started to reduce efflux. Unlike Apilimod, high levels of ML1-N was expected to sequester any unbound PI(3,5)P_2_. This is in agreement with the notion that low levels of PI(3,5)P_2_ are needed for the fusion pathway to progress, while its absence or excess have deleterious effects on the system (Miner et al., 2019).

**Figure 3.**
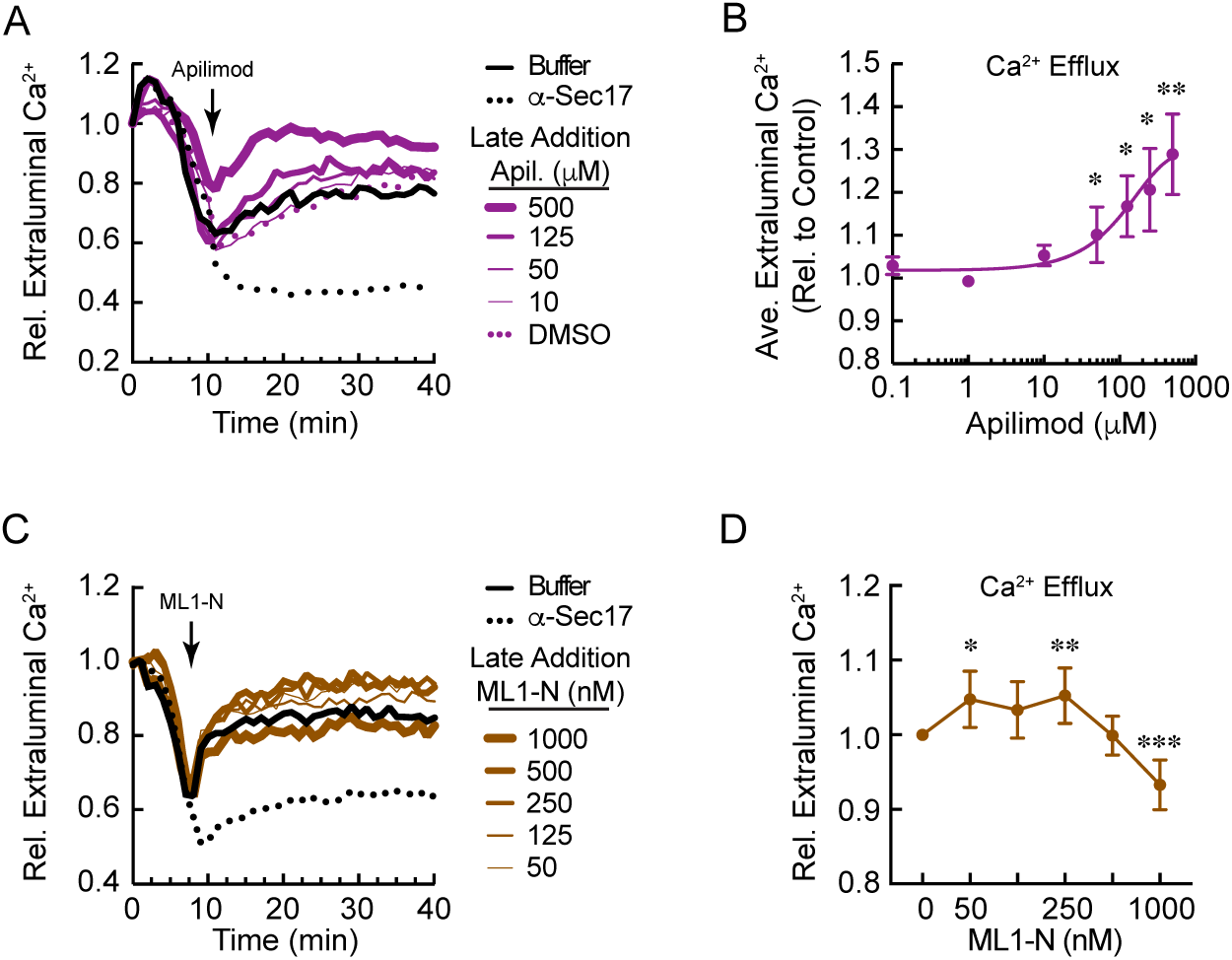
Apilimod modulates Ca^2+^ flux during vacuole fusion. (A) Vacuoles were treated with buffer of anti-Sec17 IgG and incubated for 8-10 min to allow Ca^2+^ uptake. Next, vacuoles were treated with a dose response curve of Apilimod or DMSO (Carrier). Reactions were further incubated for a total 40 min. (B) Quantitation of Ca^2+^ efflux at 30 min normalized to the untreated control. (C) Vacuoles were treated with buffer or anti-Sec17 IgG and incubated for 10 min to allow Ca^2+^ uptake. Next, vacuoles were treated with a dose response curve of GST-ML1-N and reactions were further incubated for a total of 30 min. (D) Ca^2+^ release at 30 min. Values were normalized to the untreated control at 30 min. Error bars are S.E.M. (n=3). \**p*<0.05, \*\**p*<0.01, \*\*\**p*<0.001. Significant differences were in comparison to untreated wild type efflux

### Fab1 kinase activity mutants differentially affect Ca^2+^ transport

Thus far we have used exogenous C8-PI(3,5)P_2_, chemical inhibition of Fab1, and lipid sequestration to show that Ca^2+^ efflux is affected by this lipid. To further confirm these findings, we next used vacuoles from yeast that expressed Fab1 mutations that altered kinase activity. First, we used vacuoles that contained the hyperactive Fab1 kinase mutation T2250A (Lang et al., 2017). We found that *fab1^T2250A^* vacuoles showed attenuated Ca^2+^ release early in the assay which was followed by delayed release that reached wild type levels (Fig. 4A, red). We propose that the delay in Ca^2+^ release by *fab1^T2250A^* vacuoles could be due to an increased time required for Fig4 to consume the elevated PI(3,5)P_2_ present on these organelles. This is consistent with the effects of adding exogenous C8-PI(3,5)P_2_ in Figure 1A and supports a model in which elevated PI(3,5)P_2_ concentrations suppress Ca^2+^ efflux. The delay in efflux seen with low concentrations of C8-PI(3,5)P_2_ was close to what we observed with *fab1^T2250A^* vacuoles and suggests that the prolonged inhibition by higher lipid concentrations was due to overwhelming the capacity of Fig4 to consume PI(3,5)P_2_ in the duration of the experiment. To verify that the decreased Ca^2+^ seen with *fab1^T2250A^* vacuoles was due to directly to changes in PI(3,5)P_2_ concentrations, we added ML1-N to sequester “excess” lipid and increase Ca^2+^ efflux. We found that reducing free PI(3,5)P_2_ through sequestration partially restored Ca^2+^ efflux by *fab1^T2250A^* vacuoles (Fig. 4B). Although the rescue was incomplete, the trend is consistent with the notion that elevated PI(3,5)P_2_ reduces Ca^2+^ efflux. ML1-N at concentrations at or above 500 nM reduced Ca^2+^ as seen in Figure 3. In parallel we found that Apilimod restored Ca^2+^ efflux to wild type level and continued to linearly elevated Ca^2+^ release in a dose-dependent manner (Fig. 4C)

**Figure 4.**
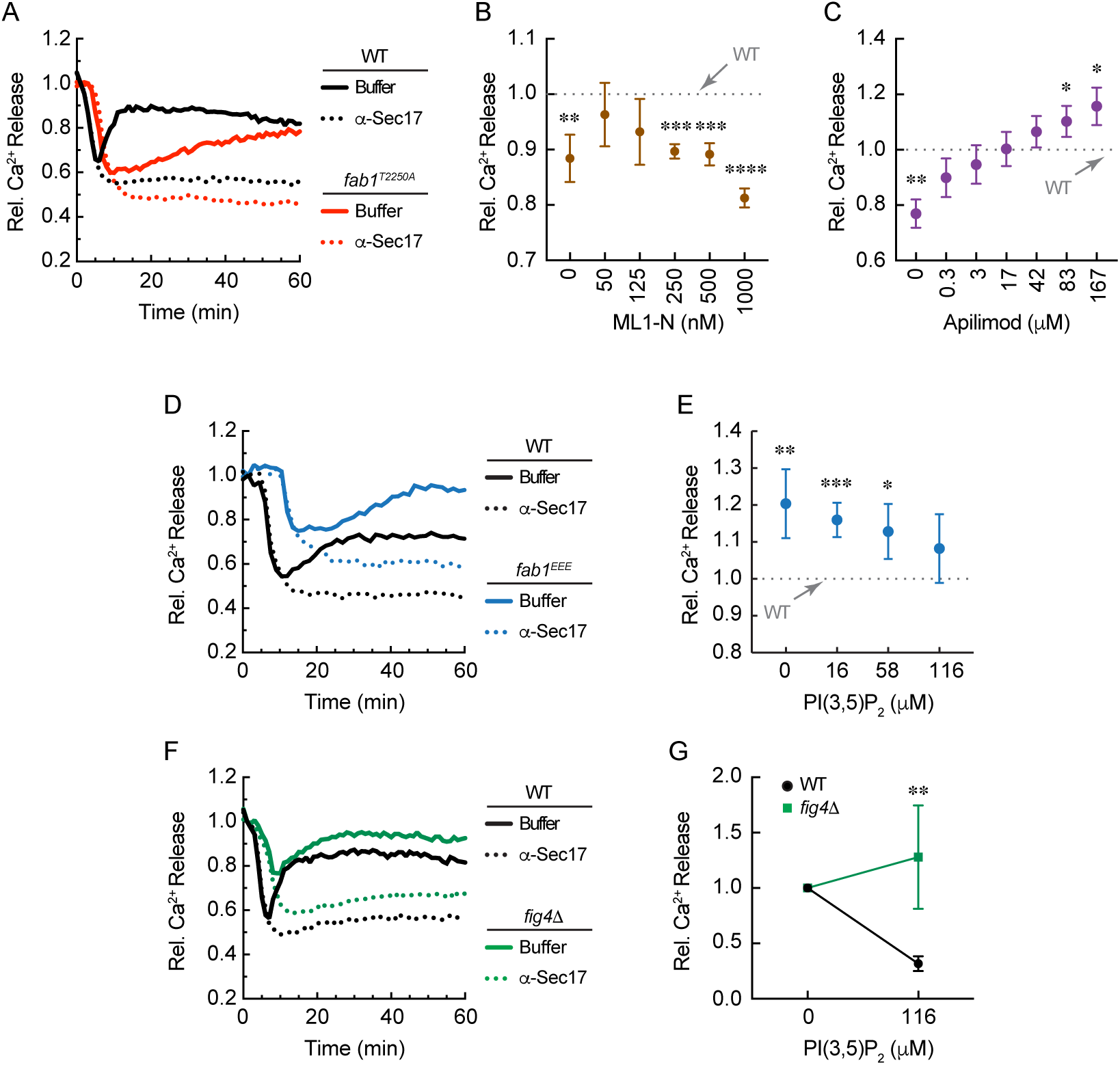
*FAB1* kinase mutations differentially affect Ca^2+^ transport. Vacuoles from wild type yeast were tested for Ca^2+^ transport in parallel with those from *fab1*^*T2250A*^ (A), *fab1*^*EEE*^ (D) or *fig4Δ* (F) yeast. (B) *Fab1*^*T2250A*^ vacuoles were treated with a dose response curve of GST-ML1-N after 10 min of incubation. Reactions were then incubated for a total of 60 min. Average Ca^2+^ efflux values at 30 min of incubation were normalized relative to wild type and given a value of 1 (dotted line). (C) *Fab1*^*T2250A*^ vacuoles were treated with a dose response curve Apilimod as described for ML1-N in (B). Values were normalized to wild type efflux at 30 min. (E) *fab1*^*EEE*^ vacuoles were treated with a dose curve of C8-PI(3,5)P_2_. Values were taken at 30 min and normalized wild type efflux without treatment at 30 min. (G) Wild type *fig4Δ* vacuoles were treated with buffer or C8-PI(3,5)P_2_. Efflux values were taken at 30 min with or without treatment for wild type and *fig4Δ* vacuoles Values were normalized to the no treatment condition for each vacuole type. Error bars are S.E.M. (n=3). Significant differences were in comparison wild type. \**p*<0.05, \*\**p*<0.01, \*\*\**p*<0.001, \*\*\*\**p*<0.0001.

To better our sense of how PI(3,5)P_2_ concentrations serve as switch to affect Ca^2+^ transport, we used vacuole from yeast that expressed the kinase dead *fab1^EEE^* mutation (Li et al., 2014). In Figure 4D, we show that *fab1^EEE^* showed a delay and overall reduction in Ca^2+^ uptake compared to wild type. The eventual release of Ca^2+^ was also delayed, yet robust. Together the altered uptake and release of Ca^2+^ by *fab1^EEE^* vacuoles reproduces the effects of adding Apilimod. To see if the effects of *fab1^EEE^* were due to the lack of PI(3,5)P_2_, we added exogenous C8-PI(3,5)P_2_ and found that the Ca^2+^ efflux of *fab1^EEE^* vacuoles was shifted to wild type levels (Fig. 4E). In contrast, our previous work showed PI(3,5)P_2_ was unable to restore the fusion defect of *fab1^EEE^* due to vacuolar maturation defects (Miner et al., 2019). This suggests that the regulation of Ca^2+^ efflux by PI(3,5)P_2_ is distinct from its role in vacuolar maturation. Collectively with the effect of overproducing PI(3,5)P_2_, these data demonstrate that PI(3,5)P_2_ can serve as a rheostat to either inhibit or enhance Ca^2+^ release from vacuoles. Furthermore, the effects of PI(3,5)P_2_ on Ca^2+^ release appear to be oppositely regulated during osmotic shock versus isotonic conditions, as elevated PI(3,5)P_2_ induces Ca^2+^ efflux under hyperosmotic conditions (Dong et al., 2010).

In parallel we also tested the effect of deleting the PI(3,5)P_2_ 5-phosphatase *FIG4* on Ca^2+^ transport. Cells lacking Fig4 have reduced levels of PI(3,5)P_2_ (Duex et al., 2006b; Duex et al., 2006a), and thus we predicted that it would have an intermediate effect compared to vacuoles with the *fab1^EEE^* mutation. As expected *fig4Δ* vacuoles showed attenuated Ca^2+^ uptake similar to *fab1^EEE^* yet did not show the same delays in Ca^2+^ uptake or efflux (Fig. 4E) Due to the reduced level of PI(3,5)P_2_ present on *fig4Δ* vacuoles we expected that Ca^2+^ efflux would be resistant to added C8-PI(3,5)P_2_. In accord with the other data, *fig4Δ* vacuoles were indeed resistant to C8-PI(3,5)P_2_ compared to wild type vacuoles (Fig. 4F). This is consistent with our previous finding with *fig4Δ* vacuoles and lipid mixing (Miner et al., 2019).

### PI(3,5)P_2_ acts to regulate Pmc1 during fusion

To hone in on the mechanism by which PI(3,5)P_2_ regulates Ca^2+^ flux during fusion we generated knockouts of the known vacuolar Ca^2+^ transporters: Pmc1, Vcx1, and Yvc1. As reported by Merz and Wickner, deletion of these transporters was unable to abolish the release of Ca^2+^ during fusion (Merz and Wickner, 2004). In keeping with their findings, we found that Ca^2+^ efflux was close to wild type with each of the deletion strains (Fig. 5A-B). We next asked if treating these deletion strains with C8-PI(3,5)P_2_ would inhibit Ca^2+^ flux to the same extent as seen with wild type vacuoles. We found that C8-PI(3,5)P_2_ inhibited Ca^2+^ efflux in all the deletion strain vacuoles tested except for *pmc1Δ* (Fig. 5A-B). Thus, it was likely that PI(3,5)P_2_ affected Ca^2+^ transport through Pmc1. Because Pmc1 transports Ca^2+^ into the vacuole, the effect on the observed efflux was likely due to the hyper-stimulation of Pmc1. It is important to emphasize that deleting *YVC1* had no effect on the inhibition of Ca^2+^ flux by PI(3,5)P_2_. While Yvc1 is essential for the PI(3,5)P_2_ activated Ca^2+^ transport during vacuole fission, this channel does not appear to play a role during vacuole fusion.

**Figure 5.**
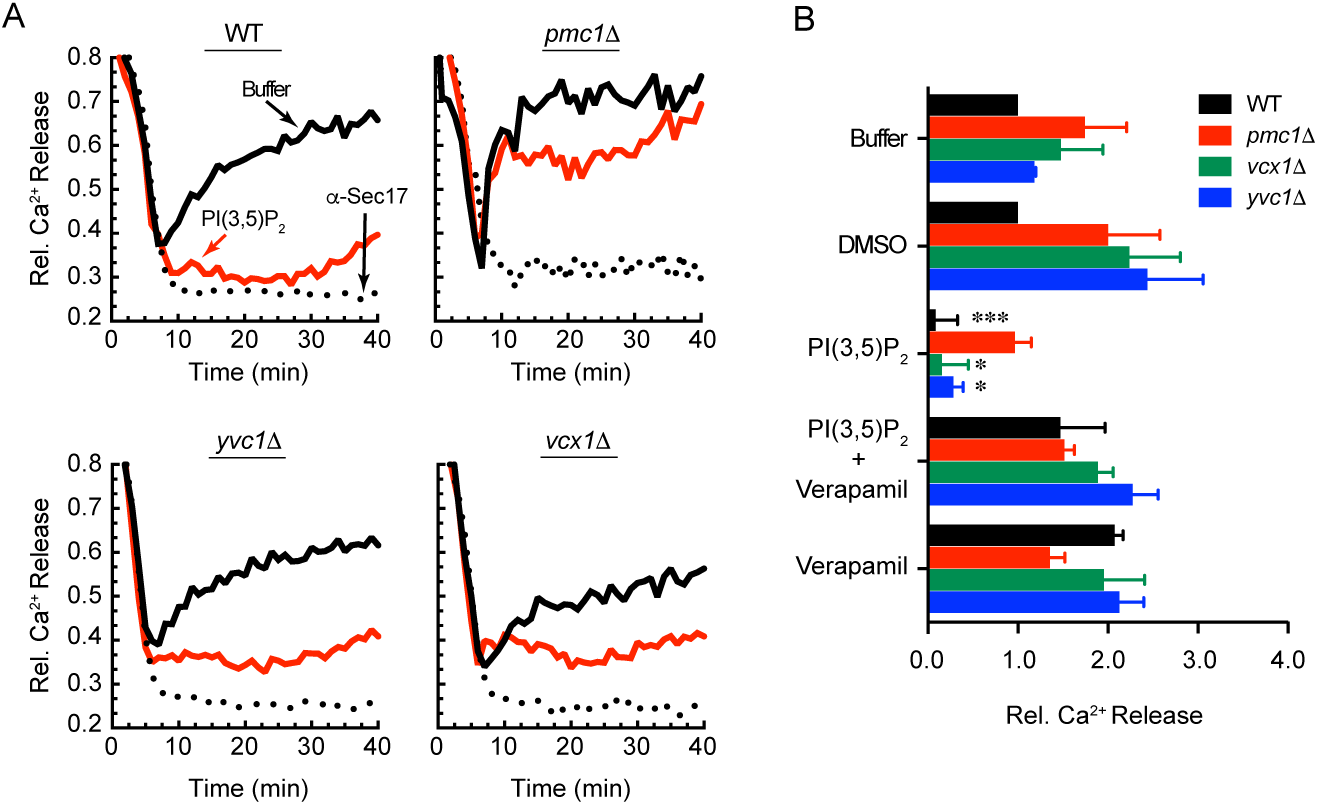
PI(3,5)P_2_ regulates Ca^2+^ transport through Pmc1 during fusion. (A) Vacuoles isolated from wild type as well as *pmc1Δ, yvc1Δ* and *vcx1Δ* strains were used in Ca^2+^ transport assays as described. Reactions were treated with anti-Sec17, buffer or 116 μM PI(3,5)P_2_ added after 5 min of incubation. Reactions were incubated for a total of 40 min and fluorescence was measured every 60 sec. (B) Average Ca^2+^ transport levels for multiple experiments with wild type, *pmc1Δ, yvc1Δ* or *vcx1Δ* vacuoles. After 10 min of incubation reactions were treated with buffer, DMSO, 116 μM C8-PI(3,5)P_2_, 500 μM Verapamil or C8-PI(3,5)P_2_ and Verapamil together and incubated for a total of 40 min. \**p*<0.05, \*\*\**p*<0.001.

If PI(3,5)P_2_ was truly acting to trigger Pmc1 activity we hypothesized that the effect should be reversible by the Ca^2+^ pump inhibitor Verapamil (Calvert and Sanders, 1995; Teng et al., 2008). In Fig 5B we summarize data showing that Verapamil blocked the effect of C8-PI(3,5)P_2_ on Ca^2+^ efflux. This further indicates that the visualized Ca^2+^ efflux could instead be the inhibition of further Ca^2+^ uptake while the cation is released at a constant rate by another mechanism.

To further test the effect of Verapamil on Ca^2+^ transport we treated vacuoles with a dose response curve of the inhibitor at the beginning of the assay. Using wild type vacuoles, we found that Verapamil inhibited Ca^2+^ uptake in a dose dependent manner when added at the beginning of the assay (Fig. 6A-B). Next, we tested if Ca^2+^ uptake continues late in the reaction by adding Verapamil after the initial uptake of Ca^2+^. As seen in Figure 6C-D, adding Verapamil resulted in a rapid accumulation of extraluminal Ca^2+^. This suggests that under control conditions, Ca^2+^ uptake continues through the docking stage, 10-15 min into the pathway, after which uptake is either inactivated or simply overcome by Ca^2+^ efflux.

**Figure 6.**
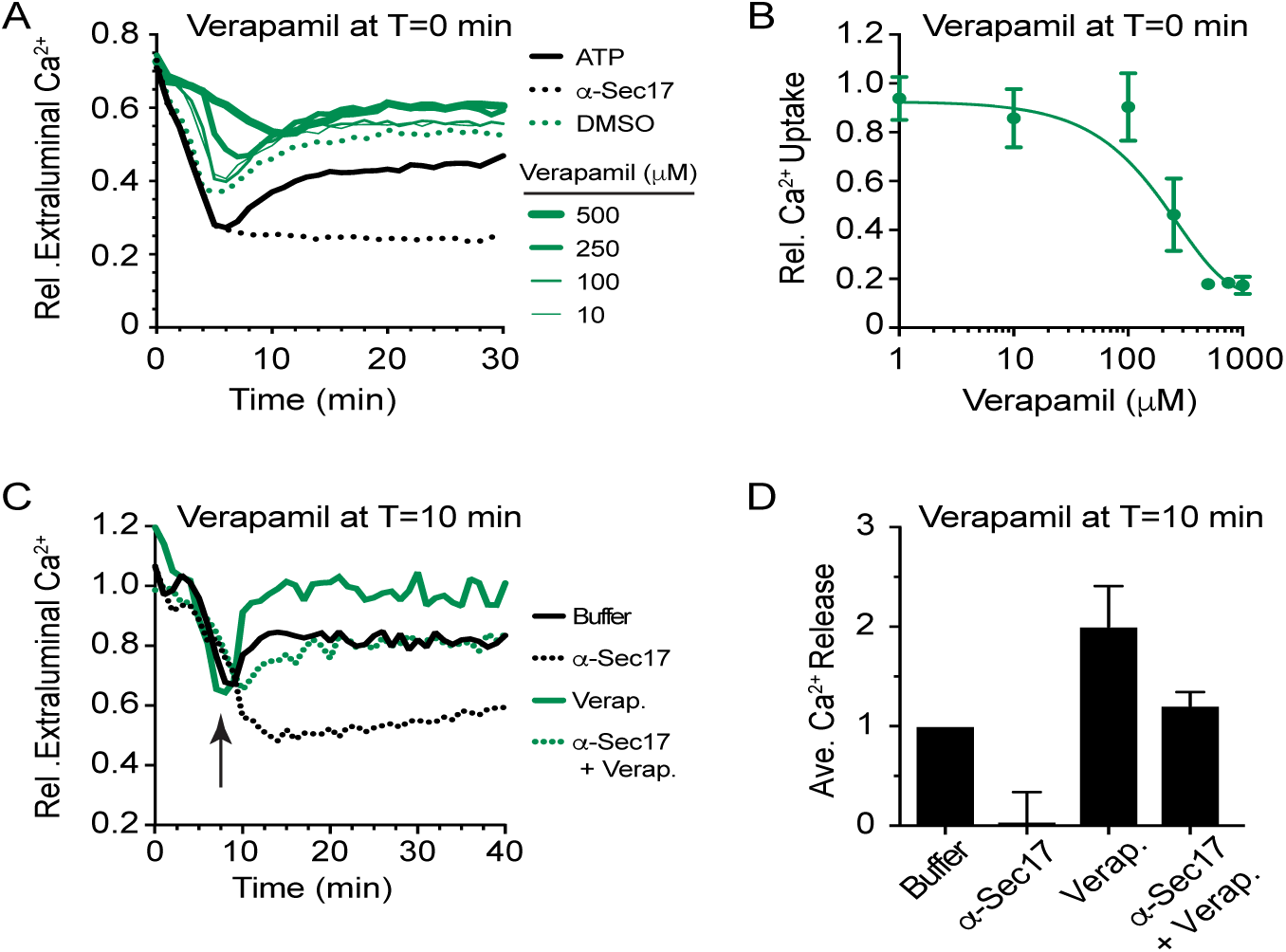
Verapamil inhibits Pmc1 activity. (A) Wild type vacuoles were treated with a dose curve of Verapamil at T=0 min to block Ca^2+^ uptake. Separate reactions were treated with buffer, DMSO or anti-Sec17 as controls. (B) Average Ca^2+^ transport levels for multiple experiments shown in panel A. Reactions were incubated for a total of 30 min and fluorescence was measured every 30 sec. (C) Wild type vacuoles were treated with 500 µM Verapamil or DMSO after 8 min of incubation to allow initial Ca^2+^ uptake. Reactions were analyzed as described in panel A. (D) Average Ca^2+^ transport levels for multiple experiments shown in panel C.

Given the surprising result that Ca^2+^ efflux appeared to be triggered by a reduction in Pmc1 activity we sought to clarify whether the main cause of Ca^2+^ efflux during membrane fusion was through the generation of transient pores as has been proposed. If transient pore formation serves as the primary efflux pathway, then we would only expect Verapamil to trigger Ca^2+^ efflux if fusion was allowed to proceed. Contrary to this explanation, we found that Verapamil was able to trigger a Ca^2+^ efflux event equivalent to standard fusion conditions even when vacuoles were treated with the membrane fusion inhibitor α-Sec17.

### PI(3,5)P_2_ regulates Pmc1 interactions with the V-ATPase

Others have previously shown that Pmc1 activity is inhibited in part through its binding to the vacuolar R-SNARE Nyv1 (Takita et al., 2001). We thus tested if added PI(3,5)P_2_ would alter the interaction. To accomplish this, we used vacuoles that harbored Pmc1 tagged with HA in order to co-immunoprecipitate Nyv1 (Cunningham and Fink, 1996). We tested Nyv1-Pmc1 interactions in the presence of buffer, C8-PI(3,5)P_2_, ML1-N and Apilimod. We found that C8-PI(3,5)P_2_ did not alter the interactions between Pmc1 and Nyv1. However, inhibiting PI(3,5)P_2_ production with Apilimod reduced the Nyv1-Pmc1 immunoprecipitation (Fig. 7A-B). Although, PI(3,5)P_2_ is not known to bind either Nyv1 or Pmc1 directly, it has been shown to bind to the V_o_ component Vph1 where it stabilizes V_o_-V_1_ holoenzyme assembly (Li et al., 2014). Because the V-ATPase is known to bind Pmc1 we tested if C8-PI(3,5)P_2_ would affect Pmc1-Vph1 interactions (Tarassov et al., 2008). Indeed, we found that PI(3,5)P_2_ reduced Vph1 interactions with Pmc1. Although, the interactions were not fully disrupted, moderate changes could have large effects on protein function. As a whole, these experiments suggest that the V-ATPase and PI(3,5)P_2_ control Pmc1 activity to modulate Ca^2+^ transport across the vacuole membrane during fusion. Although the exact mechanism by which PI(3,5)P_2_ levels alters Nyv1-Pmc1-Vph1 interactions remains to be elucidated, these findings are an exciting entrée into a new role for PI(3,5)P_2_ in vacuole homeostasis.

**Figure 7.**
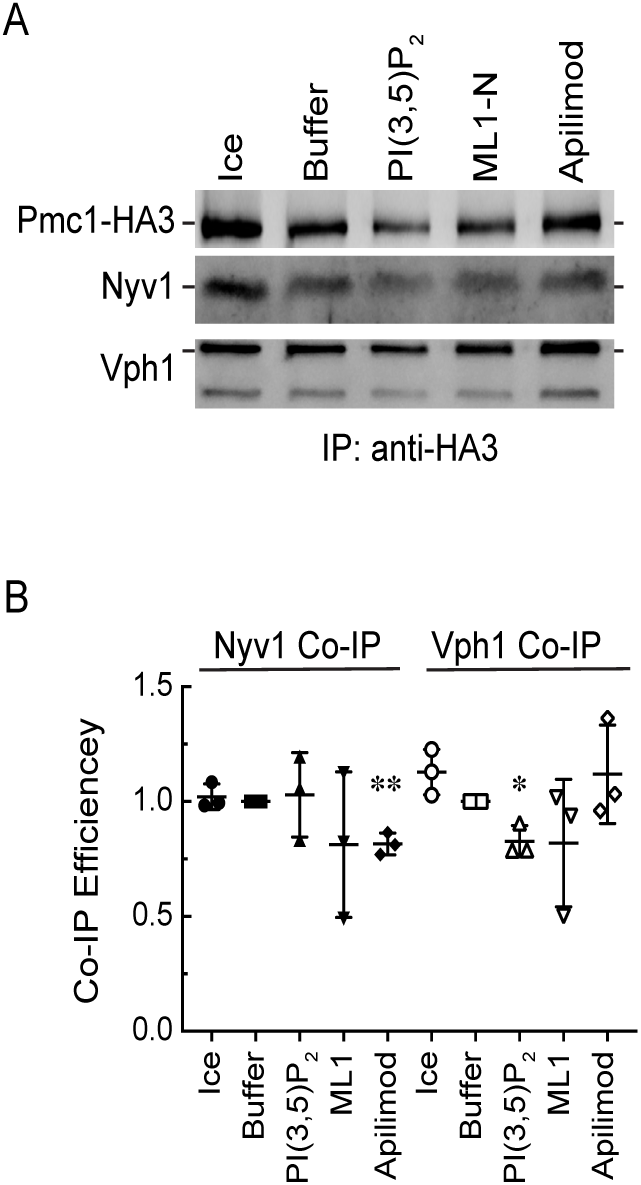
C8-PI(3,5)P_2_ enhances Pmc1 interactions with Nyv1. (A) Vacuoles expressing Pmc1-HA3 were incubated in large scale reactions (20X) in the presence of 150 µM C8-PI(3,5)P_2_, 500 nM ML1-N, 500 µM Apilimod or buffer alone. Reactions were incubated for 30 min at 27°C. After incubating, the reactions were place on ice and solubilized and processed for immunoprecipitation with anti-HA agarose. Protein complexes were eluted with sodium thiocyanate and resolved by SDS-PAGE and probed by Western blotting for the presence of Pmc1-HA3 and Nyv1. (B) Average quantitation of Nyv1 and Vph1 bound to Pmc1-HA. Values represent concentrations relative to Pmc1-HA3 at time = 30 min. Error bars represent S.E. (n=3). \**p*<0.05, \*\**p*<0.01.

## DISCUSSION

The yeast vacuole serves as the primary reservoir for Ca^2+^, which are translocated across the membrane through multiple transporters including Pmc1, Yvc1, and Vxc1. Although the regulatory mechanisms for each of these transporters are not fully understood, it has been shown that various lipids affect their function in an organelle specific manner. For instance, PI(3,5)P_2_ regulates Ca^2+^ release through Yvc1 during the osmotically driven vacuole fission (Dong et al., 2010). Ca^2+^ is also released from the vacuole during the fusion process upon the formation of *trans*-SNARE complexes (Peters and Mayer, 1998; Bayer et al., 2003; Merz and Wickner, 2004). More recently, we found that elevated levels of PI(3,5)P_2_ inhibited vacuole fusion at a stage between *trans*-SNARE pairing and outer-leaflet lipid mixing, i.e. hemifusion and full bilayer fusion (Miner et al., 2019). This was observed through adding exogenous C8-PI(3,5)P_2_ or through using *FAB1* mutations that were either kinase dead or overproduced the lipid. That study established that low levels of PI(3,5)P_2_ are needed for fusion to occur, but elevated concentrations are inhibitory. At this point it is unclear whether the effects of PI(3,5)P_2_ are direct or if the ratio with its precursor PI3P and downstream degradation product(s) also play a role. In either circumstance, it is clear that PI(3,5)P_2_ has a biphasic effect on vacuole fusion.

The biphasic nature of PI(3,5)P_2_ in vacuole fusion is not limited to this lipid. For instance, PA is needed for reconstituted proteoliposome fusion to occur and for Vam7 binding to vacuoles (Mima and Wickner, 2009; Miner et al., 2016). Yet, inhibiting PA phosphatase activity to produce diacylglycerol (DAG), resulting in elevating PA concentrations, blocks vacuole fusion at the priming stage through sequestering Sec18 from *cis*-SNARE complexes (Sasser et al., 2012a; Starr et al., 2016). Likewise, low levels of phospholipase C (PLC) stimulates vacuole fusion through converting PI(4,5)P_2_ to DAG, while high levels of PLC potently inhibits fusion (Jun et al., 2004). Finally, while PI(4,5)P_2_ is well known to stimulate vacuole fusion, an excess of the lipid can also inhibit the pathway (Fratti et al., 2004; Karunakaran and Fratti, 2013; Mayer et al., 2000; Miner et al., 2019). These examples show that the balance of substrate and product lipids can serve to move a pathway forward or arrest its progress.

In this study we continued our investigation on the inhibitory nature of PI(3,5)P_2_ on vacuole fusion. Our work has identified that elevated levels of PI(3,5)P_2_ block the net Ca^2+^ efflux observed during fusion vacuole fusion. The effect of PI(3,5)P_2_ on Ca^2+^ transport was shown by various means including the use of the kinase dead *fab1^EEE^* and hyperactive *fab1^T2250A^* mutants. While both mutants were previously shown to inhibit hemifusion (Miner et al., 2019), here we see that only *fab1^T2250A^* inhibited Ca^2+^ efflux, which is in accord with a model in which excessive PI(3,5)P_2_ is a negative regulator of the fusion pathway. In contrast, kinase dead *fab1^EEE^* vacuoles showed enhanced Ca^2+^ efflux. In both instances, Ca^2+^ flux was partially restored to wild type levels when the amount of free PI(3,5)P_2_ was increased on *fab1^EEE^* vacuoles or reduced on *fab1^T2250A^* vacuoles. The increased Ca^2+^ efflux seen with *fab1^EEE^* vacuoles observed here combined with our previous report showing that hemifusion was inhibited by this mutant indicates that Ca^2+^ flux precedes hemifusion.

Another revelation that comes from these experiments is that the effect of PI(3,5)P_2_ on Ca^2+^ transport was independent of the TRP channel Yvc1. This is the opposite to what occurs during vacuole fission induced by osmotic stress, when PI(3,5)P_2_ interacts with Yvc1 leading to the release of Ca^2+^ from the vacuole lumen (Dong et al., 2010). Our previous work with PI(3,5)P_2_ suggested that this lipid inversely regulates fission and fusion. The current work further supports this model and specifies that the effects occur through differentially modulating Ca^2+^ transport across the vacuole bilayer. Surprisingly, we found that the inhibition of Ca^2+^ transport by PI(3,5)P_2_ was dependent on the Ca^2+^ ATPase Pmc1, which utilizes ATP hydrolysis to move Ca^2+^ into the vacuole lumen against the concentration gradient. This was evident by the arrest in Ca^2+^ uptake followed by an increase in extraluminal Ca^2+^ when *pmc1Δ* vacuoles where treated with PI(3,5)P_2_. Furthermore, it appears that the regulation is immediate, as treatment with the Fab1/PIKfyve inhibitor Apilimod led to a rapid release of Ca^2+^, suggesting that PI(3,5)P_2_ needs to be newly generated, albeit at low levels to regulate Ca^2+^ transport. Under unencumbered conditions it is likely that PI(3,5)P_2_ levels are kept in check through the phosphatase activity of Fig4. One of the most striking findings from our study is that the Ca^2+^ efflux associated with fusion appears to be in part through the inactivation of Ca^2+^ influx. Additionally, our data suggests that the primary route of efflux can be uncoupled from membrane fusion as inhibition of SNARE priming was unable to prevent the trigger of Ca^2+^ efflux following treatment with Verapamil. We therefore conclude that there is a Ca^2+^ efflux route opposed by Pmc1.

The regulation of Pmc1 on yeast vacuoles is unclear relative to other P-type Ca^2+^ ATPases. Notwithstanding, others have shown that that the R-SNARE Nyv1 directly interacts with Pmc1 to reduce Ca^2+^ transport (Fig. 8) (Takita et al., 2001). Although not directly shown, their model posits that increased Nyv1 would proportionally have increased interactions with Pmc1 and reduce activity. We sought to learn if elevated PI(3,5)P_2_ levels would dissociate the Nyv1-Pmc1 complex to de-repress Ca^2+^ transport. In our hands we found that the Nyv1-Pmc1 interaction was variable when C8-PI(3,5)P_2_ was added at levels that inhibited Ca^2+^ efflux making it difficult to determine if there was an effect. However, inhibiting PI(3,5)P_2_ production with Apilimod reduced the amount of Nyv1 that co-isolated with Pmc1, suggesting PI(3,5)P_2_ can affect complex stability. This regimen of Apilimod treatment resulted in a rapid increase in extraluminal Ca^2+^. Together, these data bolster the notion that PI(3,5)P_2_ levels regulate Pmc1 activity.

**Figure 8.**
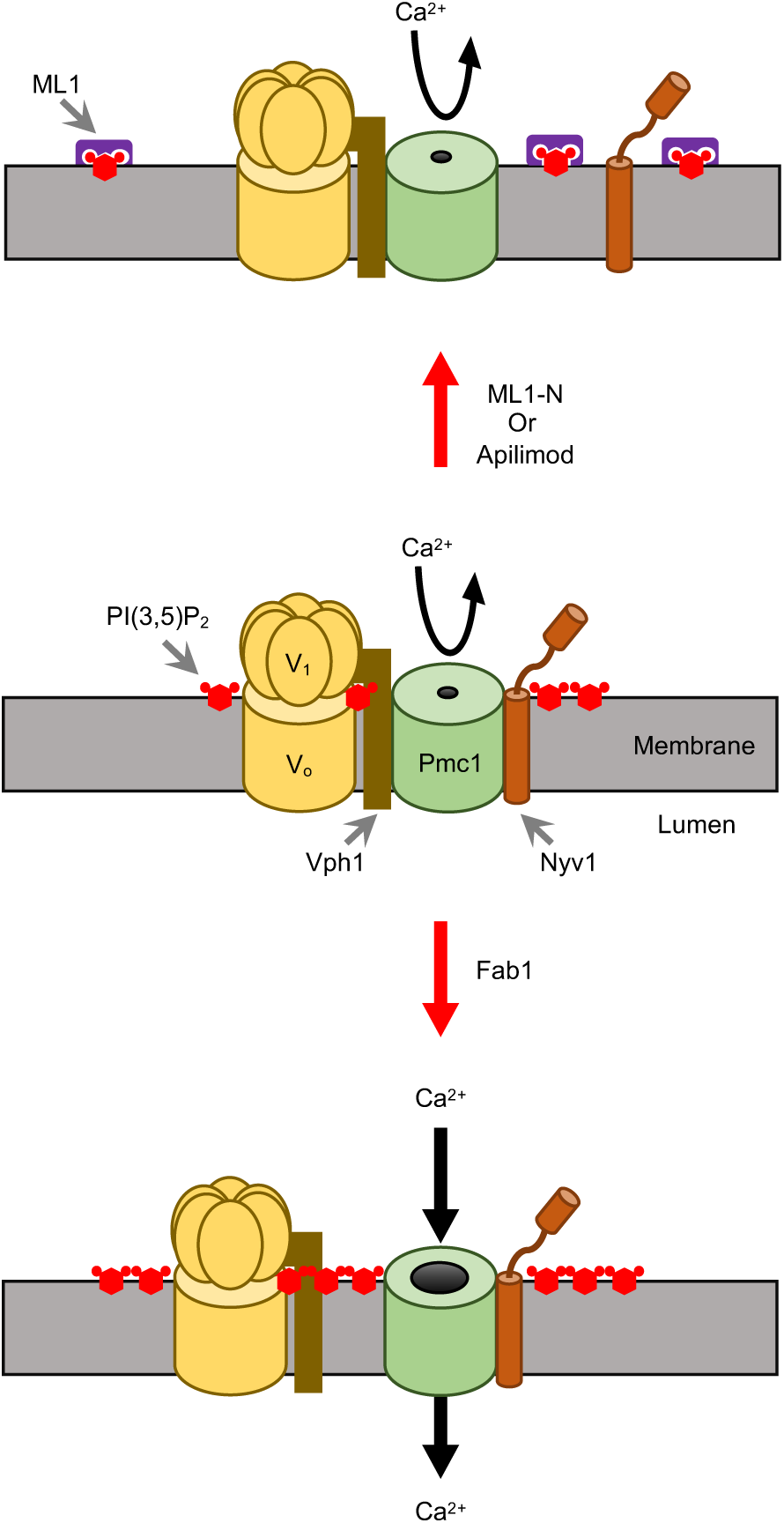
Model for PI(3,5)P_2_ regulation of Ca2+ flux during vacuole fusion.

Neither Pmc1, nor Nyv1 have been shown to bind PI(3,5)P_2_ directly, which raises the question as to how this lipid affects the complex? While a direct interaction has not been shown it is still possible that an indirect path exists. One of the few bona fide PI(3,5)P_2_ binding proteins on the vacuole is the V_o_ component Vph1 (Li et al., 2014). Their work showed that PI(3,5)P_2_ stabilized the assembly of the V_1_-V_o_ holoenzyme. Because Vph1 binds Pmc1 (Tarassov et al., 2008) and R-SNAREs have been shown to directly bind V_o_ (Di Giovanni et al., 2010), we hypothesized that PI(3,5)P_2_ could alter Vph1-Pmc1 and Pmc1-Nyv1 interactions (Fig. 8). Our Pmc1 immunoprecipitations showed that Vph1 indeed interacted with Pmc1. Furthermore, we found that adding C8-PI(3,5)P_2_ reduced the Pmc1 interaction with Vph1. When we consider that blocking V-ATPase activity results in a decrease in Ca^2+^ uptake by vacuoles (Miseta et al., 1999), we can extrapolate to think that promoting V-ATPase activity through PI(3,5)P_2_-Vph1 interactions and promotion of V_1_-V_o_ complex could inversely promote Ca^2+^ uptake by reducing Vph1-Pmc1 interactions. Our data taken together with the finding that blocking Vph1 activity through addition of *α*-Vph1 leads to inhibition of Ca^2+^ efflux (Bayer et al., 2003), we suggest a model where Vph1 binds to and inhibits Pmc1 and that the addition of C8-PI(3,5)P_2_ or α-Vph1 prevents this association. Thus, we can conclude that PI(3,5)P_2_ regulates Pmc1 in part through binding Vph1 and dissociating Pmc1-Vph1. Although this model is incomplete, we believe these data gives further insight into the mechanism behind Ca^2+^ efflux during the fusion process as well as the ability of PI(3,5)P_2_ to act as a potent fusion-fission switch.

## MATERIALS AND METHODS

### Reagents

Soluble reagents were dissolved in PIPES-Sorbitol (PS) buffer (20 mM PIPES-KOH, pH 6.8, 200 mM sorbitol) with 125 mM KCl unless indicated otherwise. Anti-Sec17 IgG (Mayer et al., 1996), and Pbi2 (Slusarewicz et al., 1997) were prepared as described previously. C8-PA (1,2-dioctanoyl-*sn*-glycero-3-phosphate), C8-PI3P (1,2-dioctanoyl-phosphatidylinositol 3-phosphate), and C8-PI(3,5)P_2_ (1,2-dioctanoyl-phosphatidylinositol 3,5-bisphosphate), BODIPY-TMR C6-PI3P and BODIPY-TMR C6-PI(3,5)P_2_ were purchased from Echelon Inc. Apilimod was from MedKoo Biosciences and Verapamil was from Cayman Chemical. Both were dissolved in DMSO as stock solutions. Anti-HA antibody was from Thermo-Fisher. GST-ML1-N was produced as described (Dong et al., 2010; Miner et al., 2019). Cy3-maleimide (GE Healthcare) was used to label GST-ML1-N according to the manufacturer’s instructions.

### Strains

Vacuoles from BJ3505 genetic backgrounds were used for Ca^2+^ flux assays (**Table S1)**. *PMC1* was deleted by homologous recombination using PCR products amplified using 5’-PMC1-KO (5’-TTCTAAAAAAAAAAAAACTGTGTGCGTAACAAAAAAAATAGACATGGAGGCCCAGAATAC –3’) and 3’-PMC1-KO (5’-TTGGTCACTTACATTTGTATAAACATATAGAGCGCGTCTACAGTATAGCGACCAGCATTC–3’) primers with homology flanking the *PMC1* coding sequence. The PCR product was transformed into yeast by standard lithium acetate methods and plated on YPD media containing G418 (250 µg/µl) to generate BJ3505 *pmc1Δ::kanMX6* (RFY84). Similarly, *VCX1* was deleted from BJ3505 by recombination using 5’-VCX1-KO (5’-TTCATCGGCTGCTGATAGCAAATAAAACAACATAGATACAGACATGGAGGCCCAGAATAC– 3’) and 3’-VCX1-KO (5’-ATATAAAAATTAGTTGCGTAAACATAATATGTATAATATACAGTATAGCGACCAGCATTC–3’) primers that flanked the *VCX1* open reading frame to make RFY86. The yeast strain expressing HA-tagged Pmc1-HA was from Dr. Kyle Cunningham (Johns Hopkins University) (Cunningham and Fink, 1996).

### Vacuole Isolation and In-vitro Ca^2+^ Flux Assay

Vacuoles were isolated as described (Haas et al., 1994). Vacuole lumen Ca^2+^ was measured as described (Miner and Fratti, 2019; Miner et al., 2016; Sasser et al., 2012b). *In vitro* Ca^2+^ transport reactions (60 µl) contained 20 µg vacuoles from BJ3505 backgrounds, fusion reaction buffer, 10 µM CoA, 283 nM Pbi2 (inhibitor of Proteinase B), and 150 nM of the Ca^2+^ probe Fluo-4 dextran conjugate MW 10,000 (Invitrogen) or Cal-520 dextran conjugate MW 10,000 (AAT Bioquest). Reaction mixtures were loaded into a black, half-volume 96-well flat-bottom plate with nonbinding surface (Corning). ATP regenerating system or buffer was added, and reactions were incubated at 27°C while Fluo-4 or Cal-520 fluorescence was monitored. Samples were analyzed using a POLARstar Omega fluorescence plate reader (BMG Labtech) with the excitation filter at 485 nm and emission filter at 520 nm for Fluo-4 or Cal-520. Reactions were initiated with the addition of ATP regenerating system following the initial measurement. The effects of inhibitors on efflux were determined by the addition of buffer, inhibitors, or C8-lipids immediately following Ca^2+^ influx. Calibration was done using buffered Ca^2+^ standards (Invitrogen).

### PI3P 5-kinase assay and Thin-Layer Chromatography

Fab1 kinase activity was measured with an assay adapted from the detection of Fig4 and Plc1 activity (Rudge et al., 2004; Jun et al., 2004) with some modifications. Kinase reactions (30 µl) contained 6 µg vacuoles from BJ3505 backgrounds, fusion reaction buffer, ATP regenerating system, 10 µM CoA, 283 nM Pbi2, 2 µM BODIPY-TMR C6-PI3P, and 1 mM sodium orthovanadate. Reaction mixtures were incubated on ice and immediately quenched with acetone (100 µl) or incubated at 27°C or on ice for 15 min in the presence of Apilimod or DMSO prior to quenching. Reactions were quenched with acetone after incubation. Following incubation acidic phospholipids were extracted from all reactions (Sasser et al., 2013). TLC plates (Partisil LK6D Silica Gel Plates (60Å), Whatman) were pretreated with 1.2 mM EGTA and 1% potassium oxalate (w/v) in MeOH/Water (3:2) and then dried at 100°C for 30 min prior to use. Dried lipids were resuspended in CHCl_3_/MeOH (1:1) (40 µl) and 5 ul was spotted on the plate. Plates were run in CHCl_3_/acetone/MeOH/AcOH/Water (46:17:15:14:8). Individual channels were loaded with PI3P and PI(3,5)P_2_ standards (Echelon). Imaging of plates was performed using a ChimiDoc MP System (BioRad) and densitometry was determined with Image Lab 4.0.1 software.

### Vacuole Docking

Docking reactions (30 µl) contained 6 µg of wild type vacuoles were incubated in docking buffer (20 mM PIPES-KOH pH 6.8, 200 mM sorbitol, 100 mM KCl, 0.5 mM MgCl_2_), ATP regenerating system (0.3 mM ATP, 0.7 mg/ml creatine kinase, 6 mM creatine phosphate), 20 μM CoA, and 283 nM Pbi2 (Fratti et al., 2004). PI(3,5)P_2_ was labeled with 2 µM Cy3-GST-ML1-N. Reactions were incubated at 27°C for 20 min. After incubating, reaction tubes were placed on ice and vacuoles were stained with 1 µM MDY-64. Reactions were next mixed with 50 μl of 0.6% low-melt agarose (in PS buffer), vortexed to disrupt non-specific clustering, and mounted on slides for observation by fluorescence microscopy. Images were acquired using a Zeiss Axio Observer Z1 inverted microscope equipped with an X-Cite 120XL light source, Plan Apochromat 63X oil objective (NA 1.4), and an AxioCam CCD camera.

### Pmc1-HA immunoprecipitation

Pmc1-HA complex isolation was performed using 20X fusion reactions. Individual reactions were treated with 150 µM C8-PI(3,5)P_2_, 500 nM GST-ML1-N, 500 μM Apilimod or buffer alone. After 30 min, reactions were placed on ice. Next, reactions were sedimented (11,000 *g*, 10 min, 4°C), and the supernatants were discarded before extracting vacuoles with solubilization buffer (SB: 50 mM Tris-HCl, pH 8.0, 2 mM EDTA, 150 mM NaCl, 0.5% Tween-20, 1 mM PMSF). Reactions were then nutated for 1 hour at 4°C. Insoluble debris was sedimented (16,000 *g*, 10 min, 4°C) and 350 µl of supernatants were removed and placed in chilled tubes. Next, 35 µl was removed from each reaction as 10% total samples, mixed with 17.5 µl of 5X SDS loading buffer and 17.5 µl PS buffer. Equilibrated anti-HA agarose beads (60 µl) were incubated with the extracts (15 h, 4°C, nutation). Beads were sedimented and washed 5X with 1 ml SB (800 *g*, 2 min, 4°C), and bound material was eluted with 0.1 M Glycine, pH 2.2 and eluates were mixed with SDS-loading buffer. Protein complexes were resolved by SDS-PAGE and examined by Western blotting.

## ACKNOWLEDGMENTS

We thank Drs. Lois Weisman (University of Michigan), Haoxing Xu (University of Michigan), and Kyle Cunningham (Johns Hopkins University) for plasmids and yeast strains. This research was supported by grants from the National Institutes of Health (R01-GM101132) and National Science Foundation (MCB 18-18310) to RAF.

## Author contributions

GEM and RAF, Conception and design, Data analysis and interpretation, Drafting or revising the article; GEM, KDS, AG, MLS, ECE BCJ Acquisition of data, data analysis

---

**Table S1.** Yeast strains used in this study

| Strain | Genotype | Source |
| --- | --- | --- |
| BJ3505 | *MATα ura3–52 trp1-Δ101 his3-Δ200* | (Jones et al., 1982) |
|  | *lys2–801 gal2 (gal3) can1 prb1-Δ1.6R pep4::HIS3* |  |
| K699 | W303-1A, *PMC1::HA* | (Cunningham and Fink, 1996) |
| RFY74 | BJ3505, *yvc1Δ::kanMX6* | (Miner et al., 2019) |
| RFY76 | BJ3505, *fab1Δ::kanMX6* | (Miner et al., 2019) |
| RFY78 | RFY76, *pRS416-FAB1<sup>T2250A</sup>* | (Miner et al., 2019) |
| RFY80 | RFY76, *pRS416-FAB1<sup>EEE</sup>* | (Miner et al., 2019) |
| RFY82 | BJ3505, *fig4Δ::kanMX6* | (Miner et al., 2019) |
| RFY84 | BJ3505, *pmc1Δ::kanMX6* | This study |
| RFY86 | BJ3505, *vcx1Δ::kanMX6* | This study |

## Abbreviations

C8: dioctanoyl
PA: phosphatidic acid
PI3P: phosphatidylinositol 3-phosphate
PI(3,5)P_2_: phosphatidylinositol 3,5-bisphospahte
SNARE: soluble N-ethylmaleimide-sensitive factor attachment protein receptor

